## Supplemental text and figures for "Partial gene suppression improves identification of cancer vulnerabilities when CRISPR-Cas9 knockout is pan-lethal"

#### Supplemental Notes

##### Relationship between RNAi efficacy and gene target expression

It has been previously reported[1] that essential genes detected using RNAi frequently have higher mRNA expression than those detected using CRISPR, a trend that we also observed among the DepMap datasets when using all high-confidence dependencies (Supplemental Fig. 10a). However, this is a surprising observation because RNAi suppression tends to have a stronger effect on cell lines with lower mRNA expression of the perturbed gene target when the target is a pan-dependency (Supplemental Fig. 10b,c), which is consistent with the detection of CYCLOPS genes using RNAi[2]. A key difference between these two different comparisons of dependency and expression level is the inclusion/exclusion of selective dependencies in the gene set being evaluated. Performing the analysis using only pan-dependencies (expected to be essential for all cell lines) is useful for estimating the technical aspects of RNAi efficacy, such as relative on-target mRNA knockdown, and shows that RNAi efficacy is stronger for targets with lower expression. Since we observe the opposite behavior using selective dependencies (RNAi efficacy is stronger for targets with high expression), it implies that subsets of cell lines are addicted to high expression of context-specific genes for which they are especially sensitive to partial inhibition despite the greater technical challenges of suppressing more highly expressed targets using RNAi.

Interpreting the meaning of the relationship between target dependency and target mRNA expression can be challenging since mRNA expression can be both an indicator of biological importance as well as a technical challenge for RNAi suppression. Therefore, we also used the protein level[3] of each target gene to determine whether it could provide additional information about whether differences between CRISPR and RNAi efficacy is biological or technical. To explore whether there are features of an essential gene target that explain whether it is detected as dependency using CRISPR or RNAi, we trained random forest classification models to distinguish the distinct CRISPR pan-dependencies from the shared CRISPR and RNAi pan-dependencies using predictive features that characterize each target gene's mRNA transcript expression or protein level[3] (Methods). We found that models using proteomics predictive features only (AUC=0.67) or proteomics and RNAseq predictive features (AUC=0.72) had better predictive accuracy than models using RNAseq predictive features alone (AUC=0.58) (Supplemental Fig. 11a). Interestingly, the correlation between protein and mRNA expression was the most important feature of the model (Supplemental Fig. 11b) where genes with higher mRNA-protein correlation (r > 0.43) across cell lines were more likely to be detected as pan-dependencies using both RNAi and CRISPR (Supplemental Fig. 11c).

**Comparisons of co-dependency networks from different experiments of the same perturbation type**

There could be complementarity between datasets of the same perturbation type as well as different perturbation types. For example, the CRISPR datasets (Avana, KY), which are typically similar to each other compared to RNAi, have some clear differences when evaluating co-dependencies. We found that top co-dependencies of selective genes were more likely to be supported by prior evidence (related genes, Methods) using the network derived from the Avana dataset, but the KY network outperforms Avana when evaluating co-dependencies of pan-dependent genes (Supplemental Figure 18a). This suggests that an experimental design parameter which differs between Avana and KY screening methods, such as screen length or cell line media, improves the discriminiation of functionally related pan-dependencies. Interestingly, there was more similar pan-dependency performance between the KY CRISPR and DRIVE RNAi networks (both 14 day screens) compared to the Avana CRISPR (21 day screen), but there are many factors besides screen length that could contribute to this result (Supplemental Figure 18c).

### Supplemental Figures

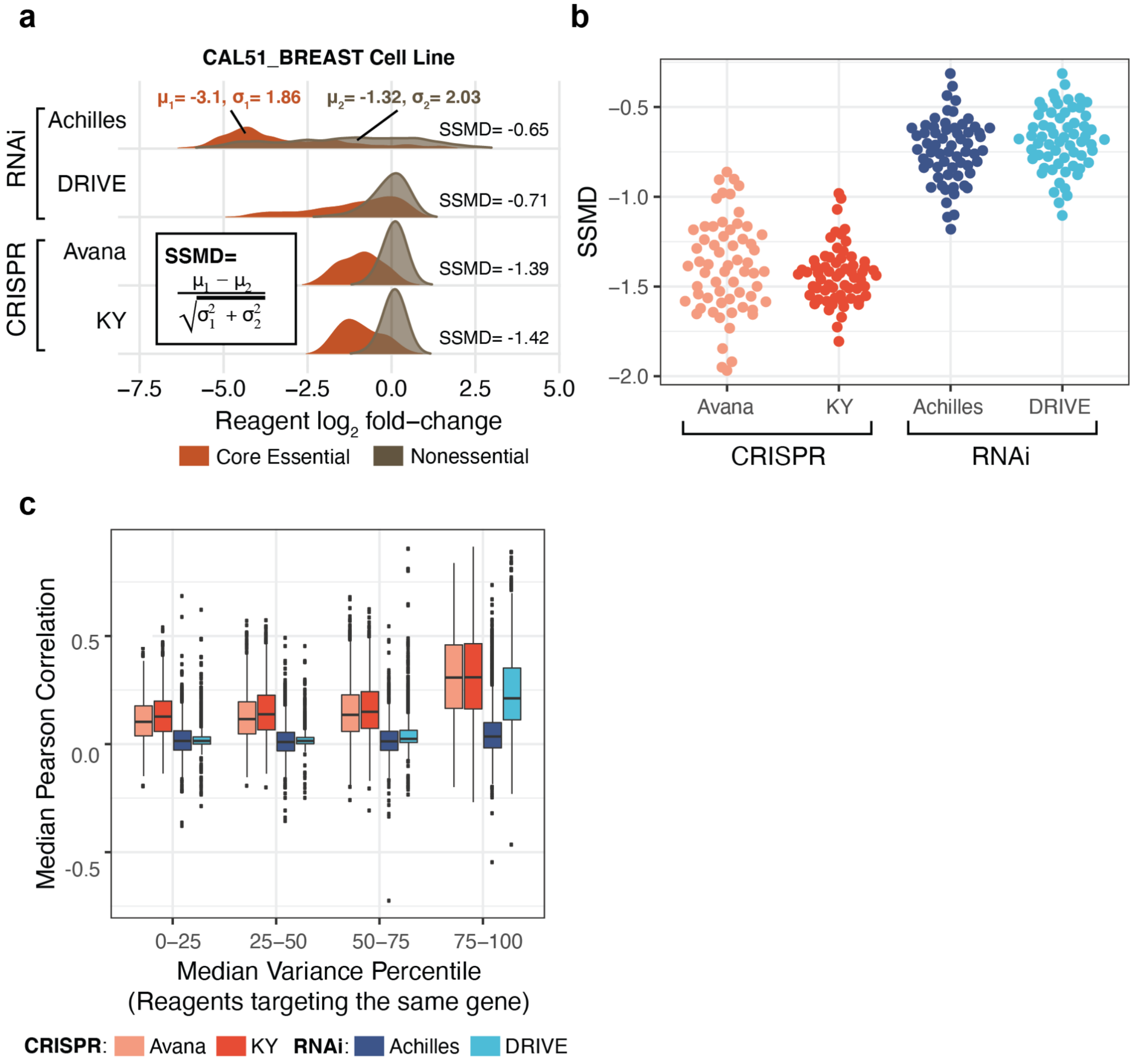


**Supplemental Fig. 1 | Quality control metrics for reagent-level screening data**

**a**, Example of the strictly standardized mean difference (SSMD) calculation for a single cell line (CAL51_BREAST), which was selected for having SSMD nearest to the average performance across all datasets. Dataset labels refer to reagent libraries (Project SCORE CRISPR: KY, DepMap CRISPR: Avana, Project DRIVE RNAi: DRIVE, Project Achilles RNAi: Achilles). SSMD represents the separation between positive (core essential[4]) and negative (nonessential[4]) controls. **b**, SSMD for cell lines (N=62) included in each reagent dataset. More negative scores indicate better screen quality. **c**, Median of Pearson correlations between all pairs of reagents targeting the same gene (y-axis). Variance per gene is based on the median variance of its corresponding reagents and represented as percentile by comparing to the median variance of all other genes within each individual dataset (x-axis). Bins are 25% of the total 7,595 genes.

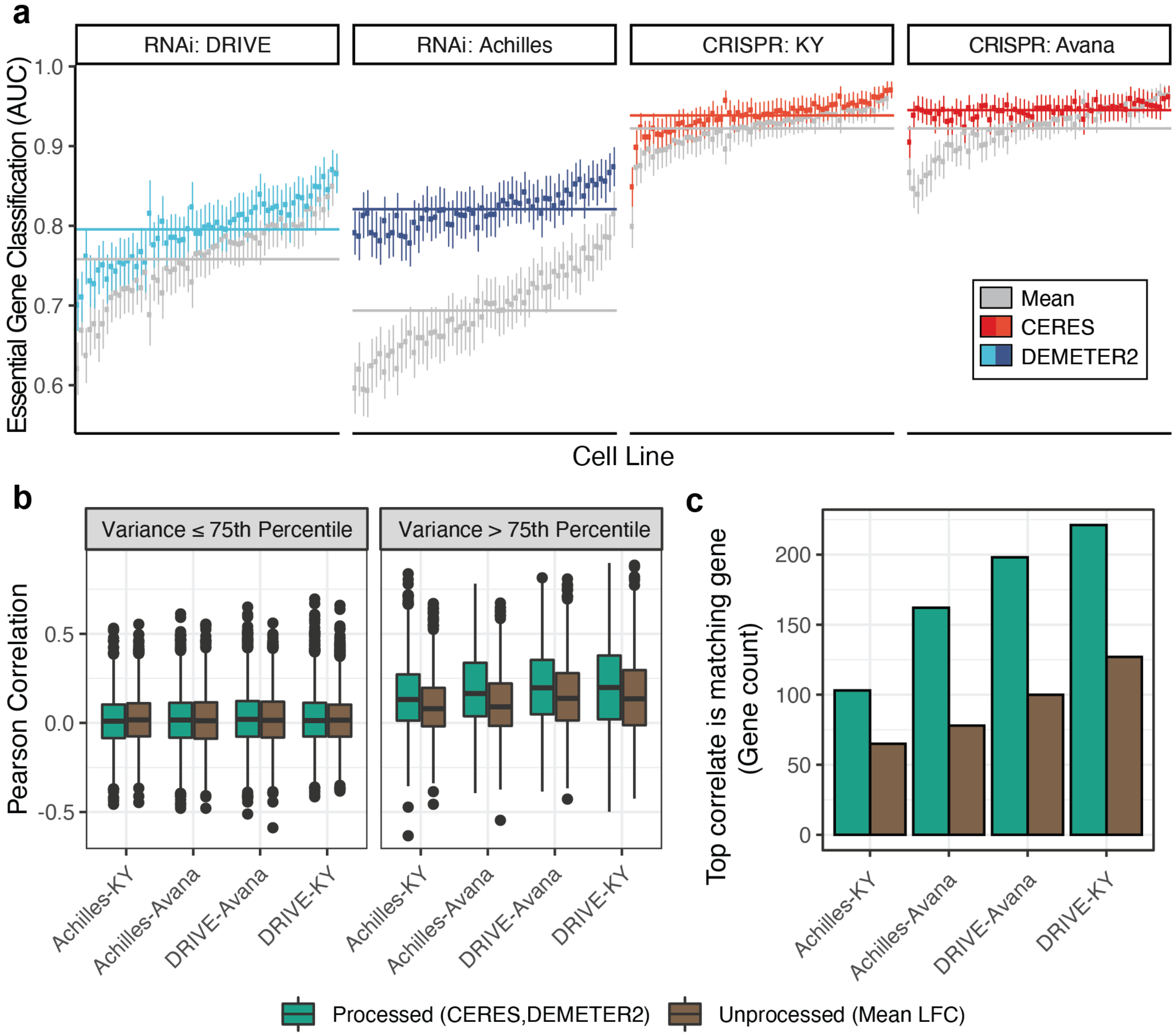


**Supplemental Fig. 2 | Comparison of processed and unprocessed data**

**a**, Receiver operating characteristic (ROC) analysis of gene effects per cell line. False positive rate and true positive rate are determined by non-expressed genes and unbiased essential genes (Methods), respectively. Values represent ROC AUC per cell line. Error bars represent 95% confidence intervals. Cell lines are aligned vertically per data source (DRIVE, Achilles, KY, Avana). Colors indicate whether the AUC values are calculated from unprocessed data (mean reagent LFC)(gray) or processed data (CERES or DEMETER2)(red,blue). Horizontal lines represent the mean ROC AUC across cell lines. **b**, Pearson correlation of matching genes between all pairwise combinations of CRISPR (Avana, KY) and RNAi (DRIVE, Achilles) datasets. Genes are binned by the mean variance of both datasets in the pair (x axis). Colors indicate processed and unprocessed data. **c**, Correlations are computed as in part b, except the correlation between matching genes is ranked in comparison to all pairwise combinations of genes between two datasets. Bars represent the number of genes (out of 5,226 total genes shared between all datasets) for which the matching gene is the top correlate.

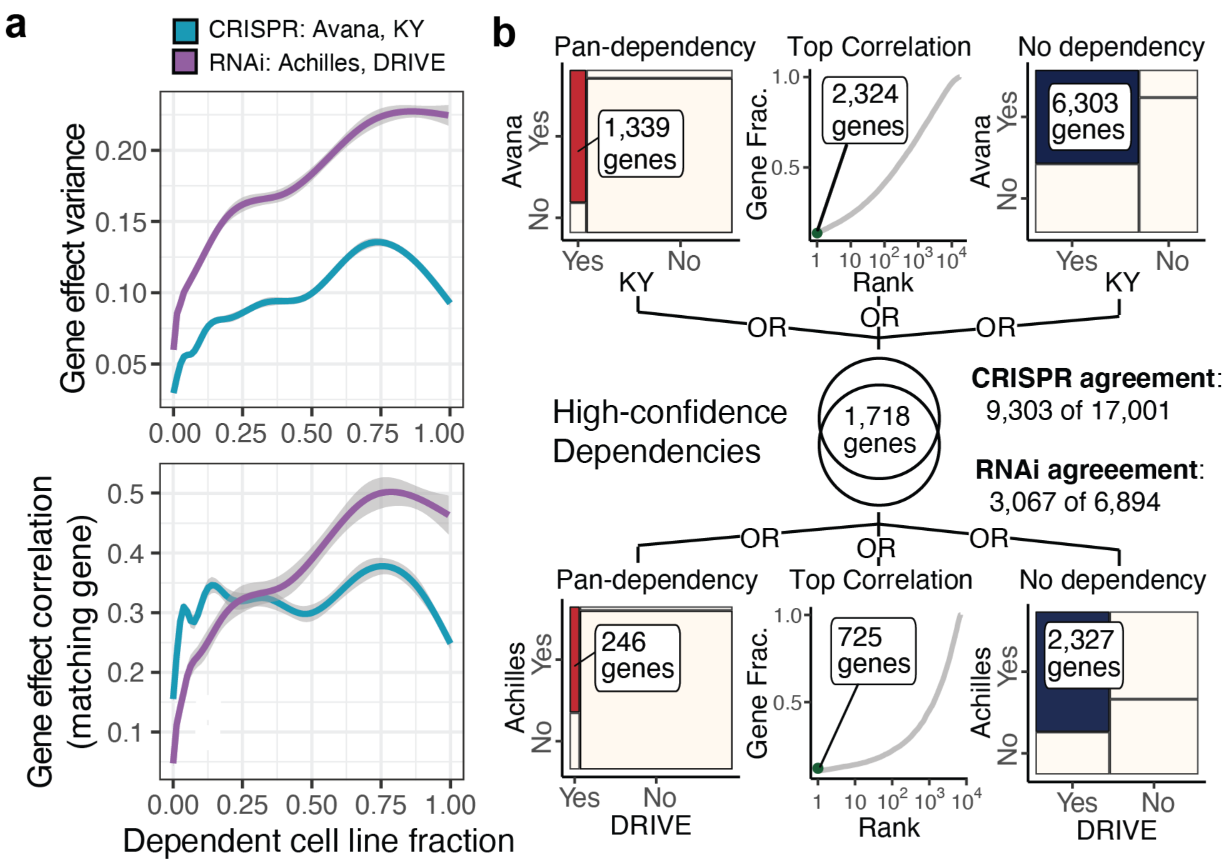


**Supplemental Fig. 3 | Identifying high-confidence dependency profiles that agree between gene effect datasets from the same perturbation type**

**a**, The RNAi datasets are the DEMETER2 processed Project Achilles and Project DRIVE datasets. The CRISPR datasets are the CERES processed DepMap ‘Avana’ dataset and the Project SCORE ‘KY’ dataset. The x axis represents the fraction of dependent cell lines (probability of dependency > 0.5) per gene, calculated per dataset and averaged according to perturbation modality (CRISPR, RNAi). The gene effect variance across cell lines (top) and gene effect correlation between datasets (bottom) are shown as a function of the gene dependency fraction. The gene effect variance is normalized per dataset and mean collapsed by perturbation modality (CRISPR, RNAi). **b**, Criteria for inclusion in the high confidence dependency set. Genes that are classified as pan-dependencies or non-dependencies by the respective analyses in both library’s datasets, or genes that are top ranking correlations (as shown in part b), are considered to be in agreement across libraries. The CRISPR libraries have 9,303 genes with multi-library agreement out of 17,001 shared targets (3,067 out of 6,894 shared targets for RNAi). The intersection of CRISPR and RNAi multi-library agreement gene sets results in 1,718 high-confidence CRISPR and RNAi dependency profiles. Of these 1,718 high-confidence dependencies, 1,703 are included in the 15,221 genes that overlap the DEMETER2-Combined RNAi and DepMap CRISPR dataset.

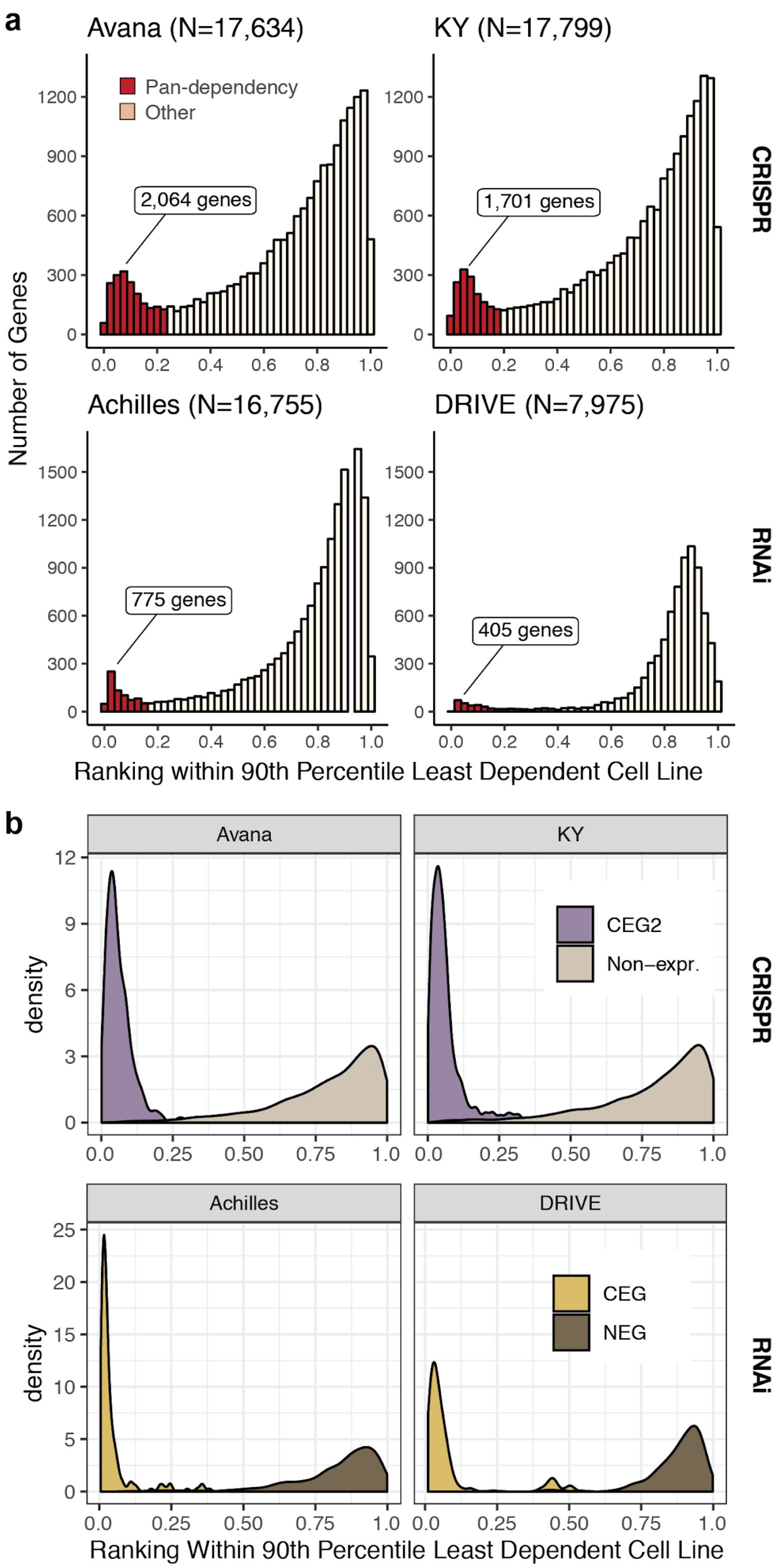


**Supplemental Fig. 4 | Method of identifying pan-dependencies in each dataset**

**a**, Genes are ranked per cell line by ascending CERES (CRISPR) or DEMETER2 (RNAi) gene effect and normalized from 0 to 1. For each gene, the normalized rank of the 90th percentile least dependent cell line is used as a pan-dependency score. The histograms represent the pan-dependency scores for all genes in each dataset. Genes in the left-most mode (red) with labeled count totals are classified as pan-dependencies. **b**, Distribution of the same metric used to classify pan-dependencies in (**a**) for all genes included in benchmark essential (RNAi: Core essential genes[4], CRISPR: Expanded core essential genes[5]) and non-essential (RNAi: Nonessential genes[4], CRISPR: Non-expressed [TPM < 0.2 in greater than 50% of cell lines]) gene sets.

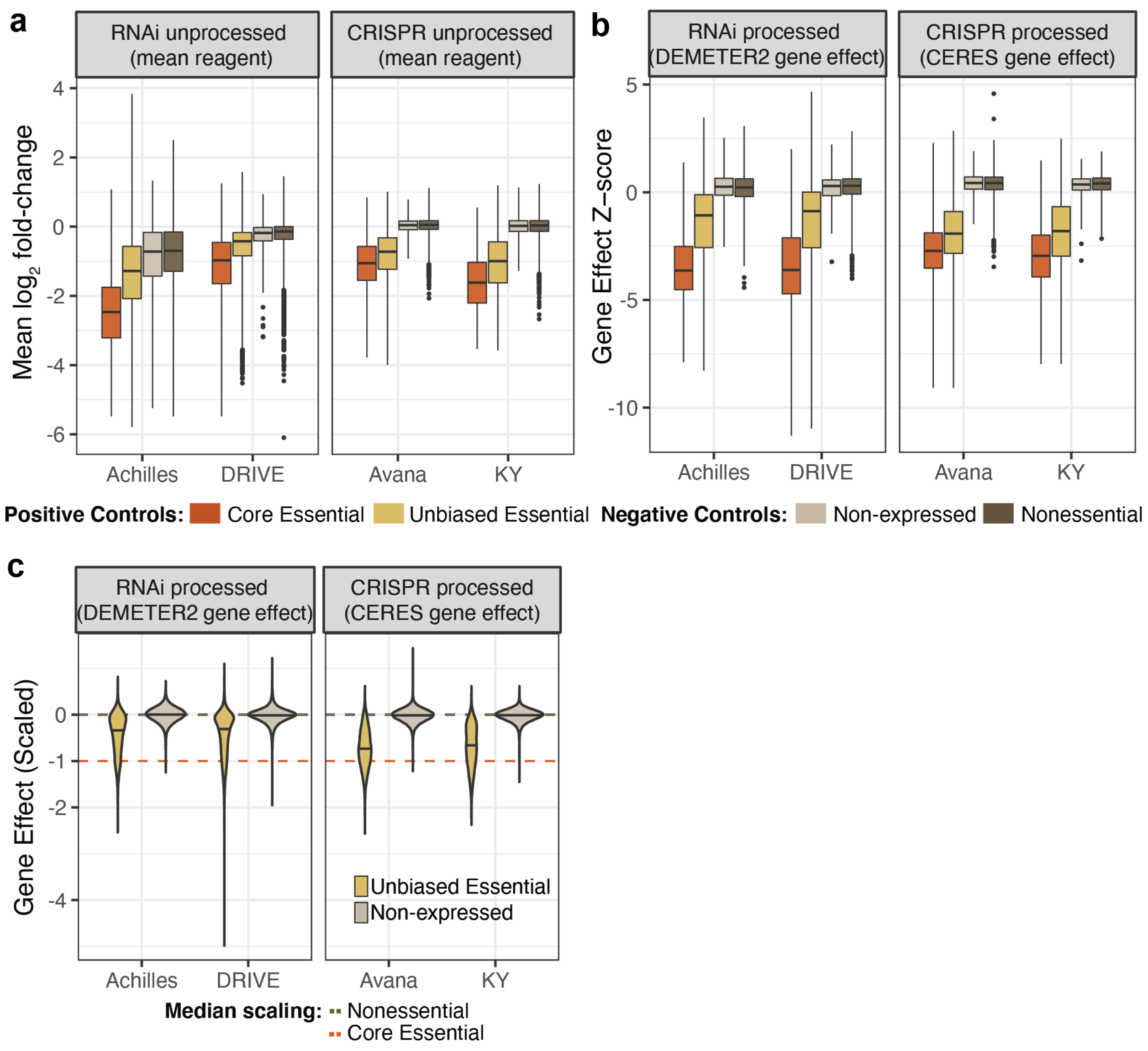


**Supplemental Fig. 5 | Efficacy and specificity estimates using control sets**

**a**, Reagent $\log_2$ fold-change (LFC) datasets are filtered for the 7,595 gene targets and 62 cell lines that overlap all four datasets. LFC values for reagents targeting the same gene are collapsed by a simple average (y-axis) per cell line. Boxes represent the distribution of mean LFC values for all cell lines that target genes included in the nonessential genes[4], non-expressed set (10 randomly sampled genes from each cell line with RNAseq $\log_2$(TPM+1) < .2), unbiased essential gene set described in the Methods, or the core essential genes[4]. **b**, Gene effect estimates from DEMETER2 (RNAi) or CERES (CRISPR) for 6,714 shared gene targets were Z-scored per cell line (N=69). Boxes represent all gene effect z-scores from all cell lines that are included in the corresponding benchmark gene sets from part *a*. **c**, Processed gene effects from part *b* are scaled per cell line using the standardized method from the Project Achilles pipeline (medians of the nonessential genes are 0 and the median of the core essential genes are -1). Violin plots show the distribution of genes included in control sets (unbiased essential genes, non-expressed genes) that were not used for scaling. Horizontal lines represent the median of core essential and nonessential genes per cell line.

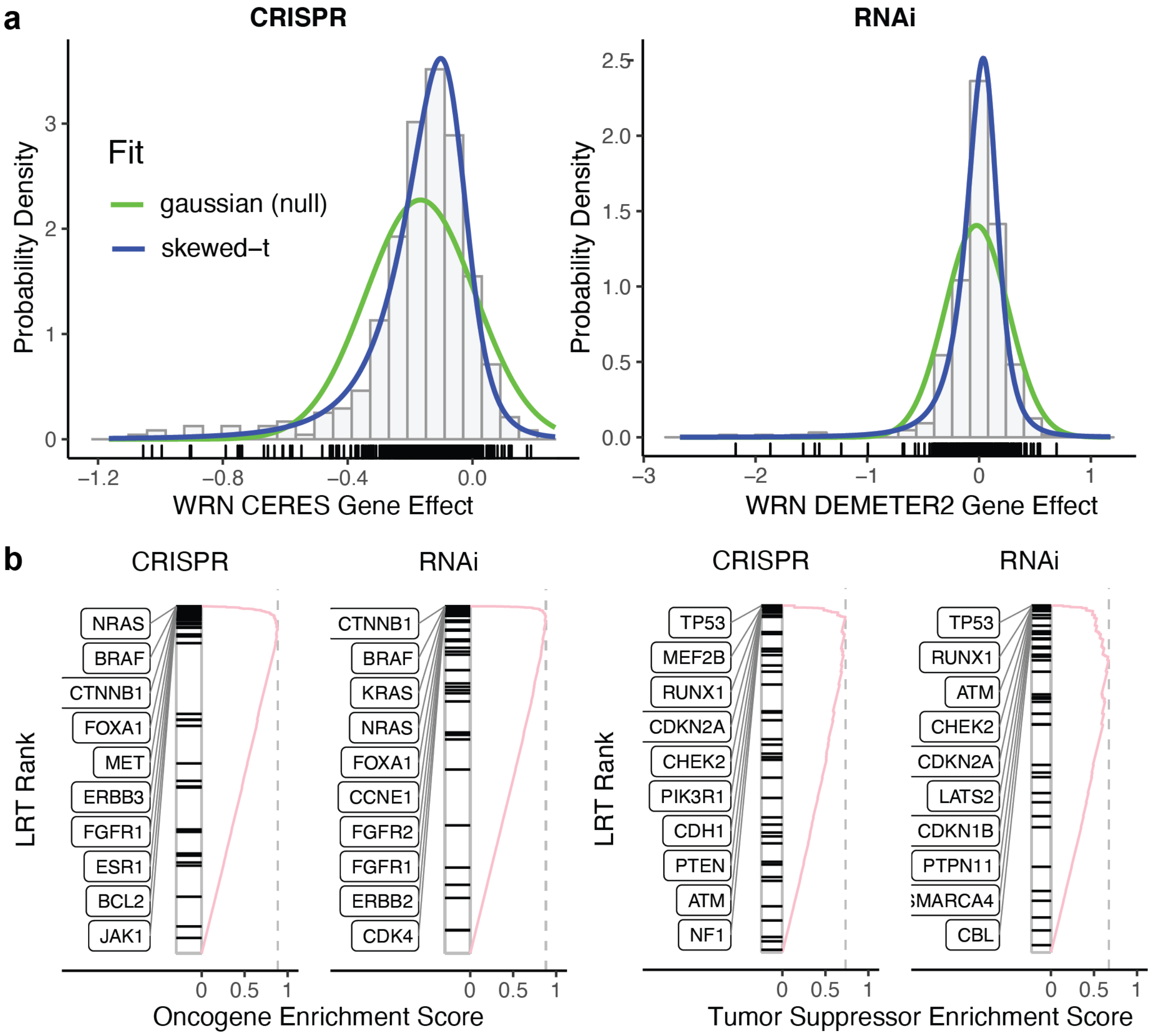


**Supplemental Fig. 6 | Method of identifying strongly selective dependencies**

**a**, Example of likelihood-ratio test (LRT) calculation for WRN dependency using CRISPR and RNAi gene effects. The LRT score for WRN is the ratio of the likelihoods that the observed WRN gene effects (grey bars) fit a skewed-t (blue) distribution compared to the fit to a gaussian (green) distribution. A higher LRT score indicates that the gene effects are more likely under the skewed-t than the gaussian null. We define a strongly selective dependency (SSD) to be a gene effect profile with LRT score > 100. WRN qualifies as an SSD using both CRISPR or RNAi gene effects. **b**, Significant enrichment for oncogenes (CRISPR and RNAi p values = $1 \times 10^{-6}$) and tumor suppressors (CRISPR p value = $8 \times 10^{-6}$, RNAi p value = $1.73 \times 10^{-4}$) among genes with high LRT scores. Oncogenes and tumor suppressor genes (TSGs) are downloaded from the Oncology Knowledge Base (OncoKB). Oncogenes are filtered for those with at least one hotspot mutation (frequent occurrence in TCGA or COSMIC) in the set of 403 cell lines being compared between RNAi and CRISPR, resulting in 58 oncogenes. Similarly, the TSGs are filtered for genes where at least one cell line has a mutation that is predicted to be deleterious, resulting in 46 TSGs. Statistical significance for the enrichment of oncogenes and TSGs among the top ranked LRT scores for CRISPR (CERES Avana) and RNAi (D2-Combined) are computed using GSEA with default weight parameter of 1 and a million permutations.

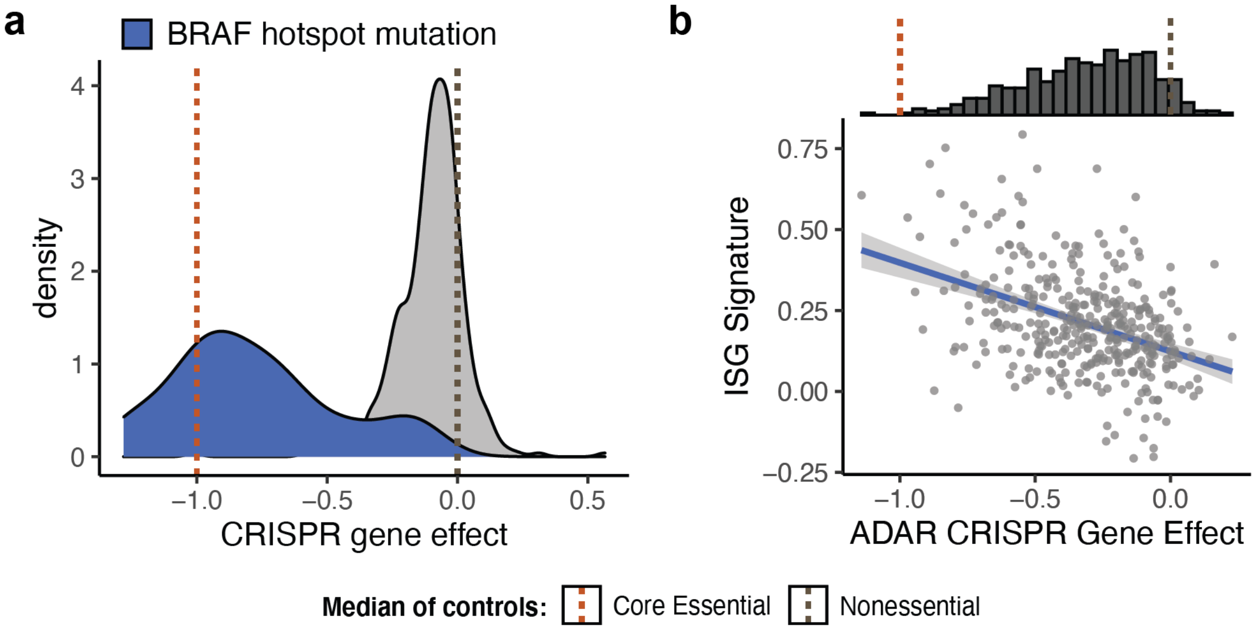


**Supplemental Fig. 7 | Examples of selective dependencies**

**a**, Distribution of BRAF gene effect (CRISPR) across cell lines, colored by presence (blue) of absence (gray) of a BRAF mutation (identified as a hotspot mutation by TCGA or COSMIC). This type of dependency distribution has a high skewed-t to gaussian LRT score and is classified as strongly selective. **b**, The distribution of ADAR gene effects are not as skewed as BRAF, making it not as easily identified as a selective dependency using the LRT score. Although there isn't a clear subset of dependent cell lines, the gene effects are correlated with an established predictor of ADAR dependency, interferon-stimulated gene (ISG) expression signature (Methods), suggesting that this type of high-variance selectivity could also indicate potential cancer vulnerability. Instead, we use the variance in probability of dependency beyond the 99th percentile of the non-expressed genes to identify this type of selectivity.

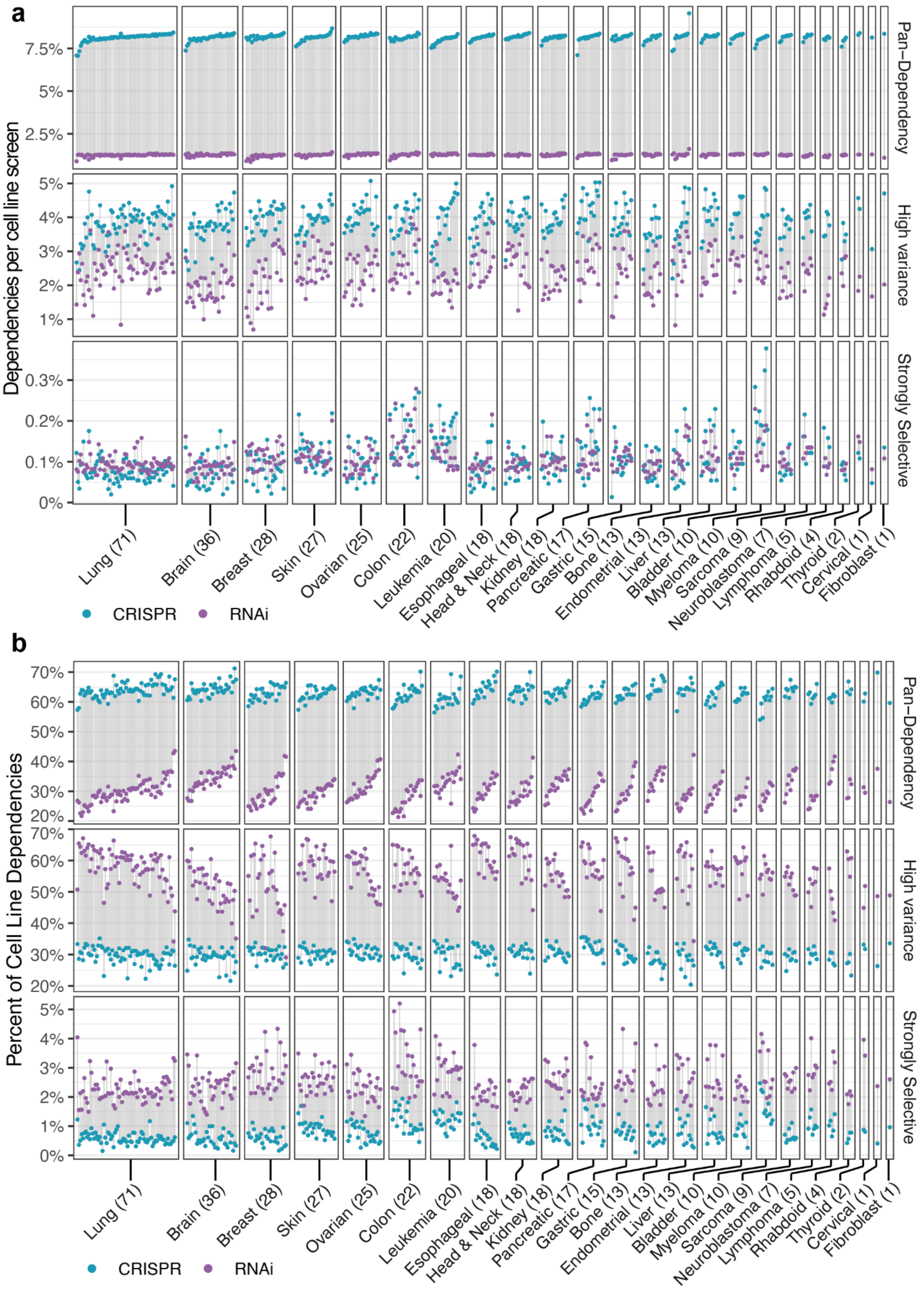


**Supplemental Fig. 8 | CRISPR knockout identifies a higher proportion of pan-dependent genes than RNAi knockdown across cell lines of all tissue origins**

**a**, Cell lines screened using both CRISPR and RNAi (N=403) are aligned vertically (connected by grey lines) and grouped by disease type (sample sizes in parentheses). The y-axis value represents the percent of genes that are dependencies in each cell line with respect to the total number of genes profiled with both technologies. The vertical facets separate genes classified as pan-dependencies, high-variance dependencies, or strongly selective dependencies (SSDs) by CRISPR (blue) or RNAi (purple). **b**, Same as part a, except the y-axis represents the percent of gene dependencies per cell line instead of the percent of total genes profiled.

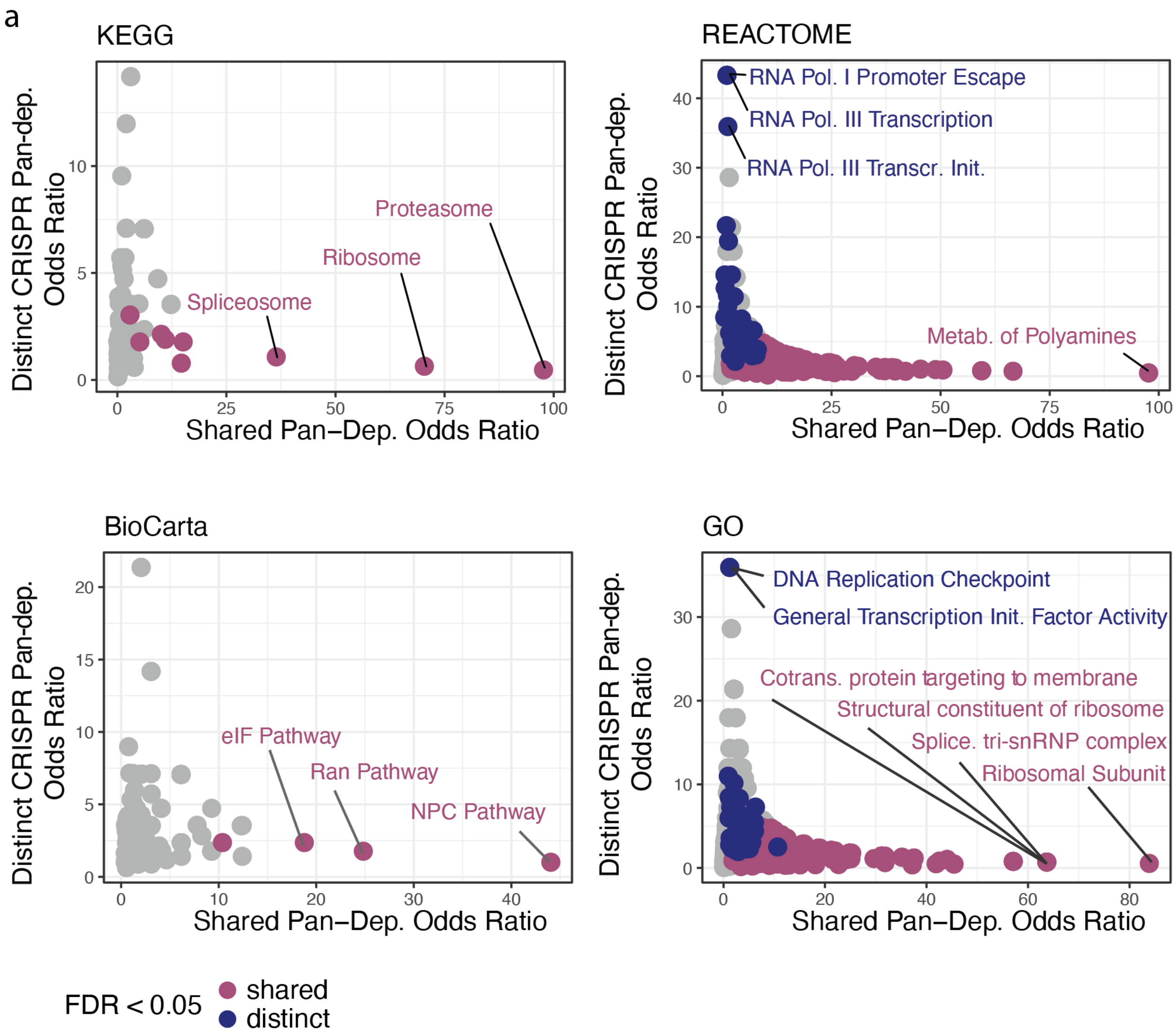


**Supplemental Fig. 9 | Gene set enrichment of distinct versus shared pan-dependencies**

**a,** Gene set enrichment of high confidence pan-dependencies that are shared (N=234) by RNAi and CRISPR or distinct (N=208) to CRISPR. Odds ratio and P-values produced by Fisher's exact test of 2-by-2 contingency table where the in-group is either shared or distinct pan-dependencies and the out-group is all other high confidence dependencies. Colored gene sets have a FDR < 0.05 where FDR is an adjusted p-value for multiple hypothesis testing across all gene set collections (N=12,216). Significant sets (FDR < 0.05) are colored (blue, purple). Gene sets include Biocarta (N=289), GO (biological process, cellular component, molecular function) gene sets (N=9,996), KEGG (N=186), and Reactome (N=1,499).

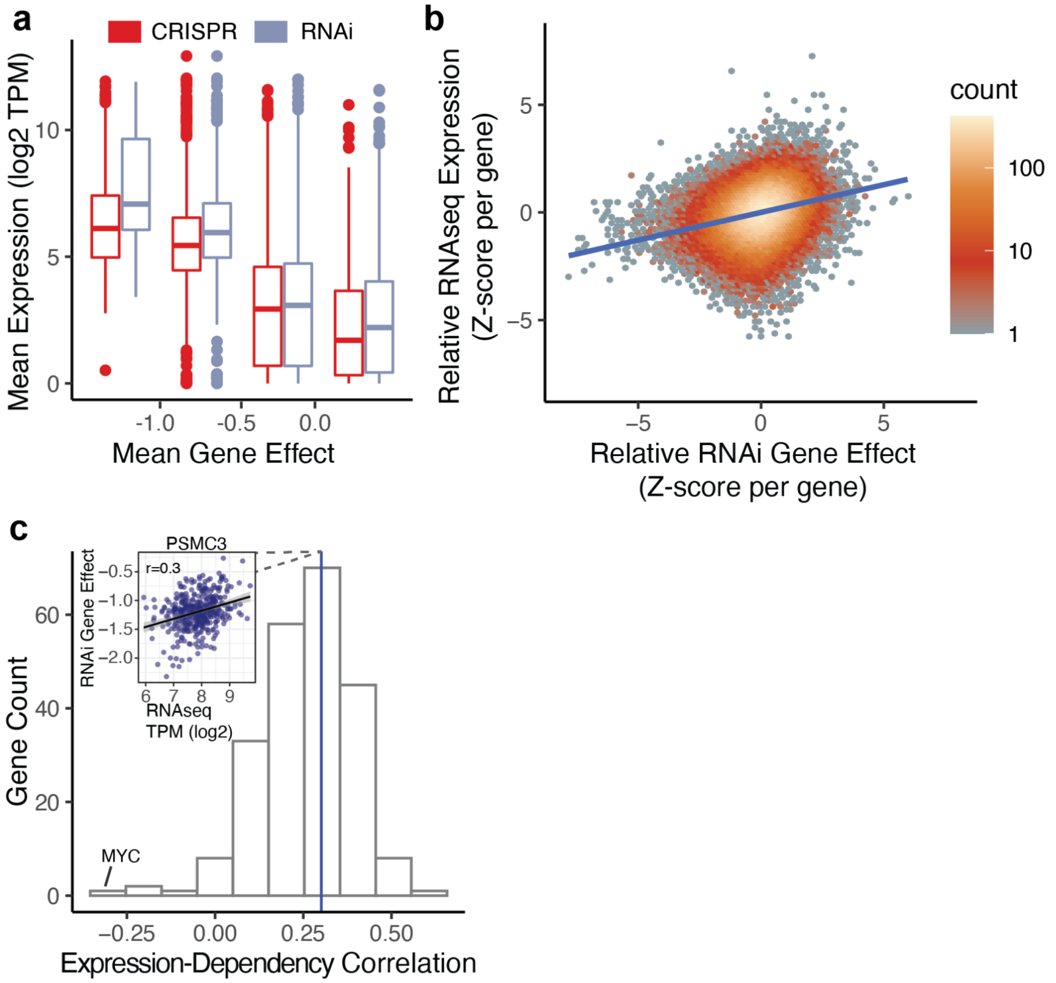


**Supplemental Fig. 10 | Association between mRNA expression and RNAi gene effect**

**a**, For each high-confidence dependency (N=1,716), including selective dependencies, boxes represent the mean RNAseq expression (TPM) across cell lines, binned by mean CRISPR (CERES Avana) or RNAi (D2-Combined) gene effect. All boxes represent over 150 genes. **b**, Using all RNAi pan-dependencies filtered for high-confidence, the RNAi gene effects and RNAseq expression profiles are Z-scored per gene and then aggregated together. Z-score values are plotted for all cell lines across 226 high-confidence RNAi pan-dependencies and are colored by the number of values in the 2D area of the point. **c**, Histogram of Pearson correlation coefficients between RNAseq expression (log TPM) and RNAi D2-Combined gene effect for the 226 RNAi pan-dependencies from part (a). Scatter plot shows an example of the underlying data for the correlation coefficient of a single gene, PSMC3. While most pan-dependencies have a positive correlation between expression and RNAi gene effect (cell lines with lower expression tend to exhibit decreased cell fitness after suppression with RNAi), there are negatively correlated outliers, such as MYC, which is a rare combination of pan-dependency and oncogene.

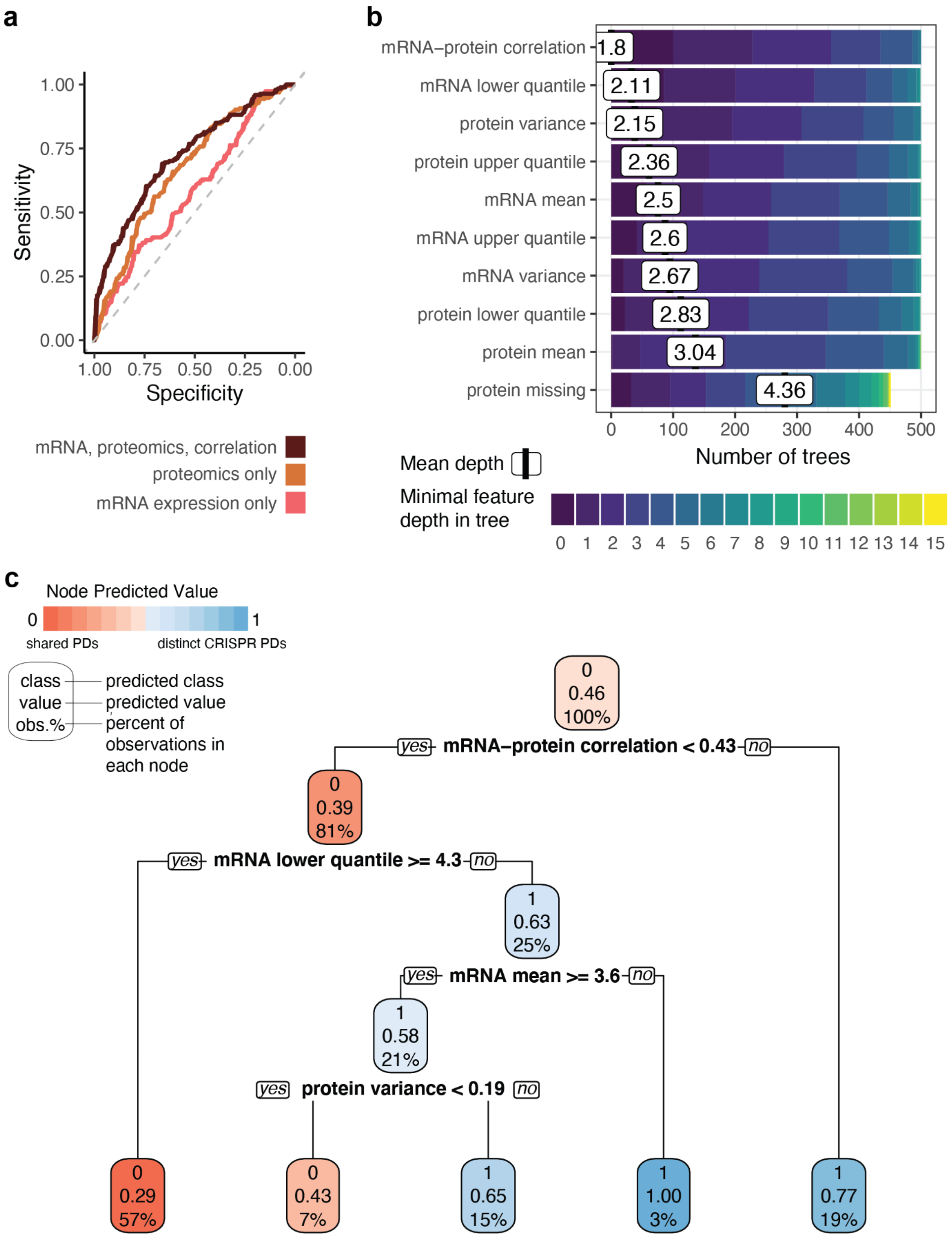
a
1.00
0.75
0.50
0.25
0.00
Sensitivity
1.00 0.75 0.50 0.25 0.00
Specificity
mRNA, proteomics, correlation
proteomics only
mRNA expression only
b
mRNA−protein correlation 1.8
mRNA lower quantile 2.11
protein variance 2.15
protein upper quantile 2.36
mRNA mean 2.5
mRNA upper quantile 2.6
mRNA variance 2.67
protein lower quantile 2.83
protein mean 3.04
protein missing 4.36
0 100 200 300 400 500
Number of trees
Mean depth
Minimal feature depth in tree
0 1 2 3 4 5 6 7 8 9 10 11 12 13 14 15
c
Node Predicted Value
0
1
shared PDs
distinct CRISPR PDs
class — predicted class
value — predicted value
obs.% — percent of observations in each node
0 0.46 100%
yes mRNA−protein correlation < 0.43 no
0 0.39 81%
yes mRNA lower quantile >= 4.3 no
1 0.63 25%
yes mRNA mean >= 3.6 no
1 0.58 21%
yes protein variance < 0.19 no
0 0.29 57%
0 0.43 7%
1 0.65 15%
1 1.00 3%
1 0.77 19%

**Supplemental Fig. 11 | Predicting distinct CRISPR pan-dependencies from mRNA and proteomics features of the target gene**

**a**, Multivariate classification accuracy of pan-dependencies that were detected using both CRISPR and RNAi datasets from those that are distinct to the CRISPR dataset. Two different random forest models were trained using predictive features (Methods) that describe either the distribution of the perturbed gene's protein level (ROC AUC=0.68) or mRNA expression level (ROC AUC=0.6). A third model uses all features (ROC AUC=0.72), including both protein and mRNA features as well as the Pearson correlation between mRNA and protein profiles for the perturbed gene. **b**, Minimum depth for the top 10 predictive features across 500 trees from the random forest model trained using all predictive features from part *a*. Lower values indicate the feature is used closer to the root of the tree (root = 0) and imply greater feature importance. **c**, Single tree fit using all pan-dependencies, but only the top 5 features according to mean minimal depth from part *b* were used for a simplified example of how these features interact within a single tree.

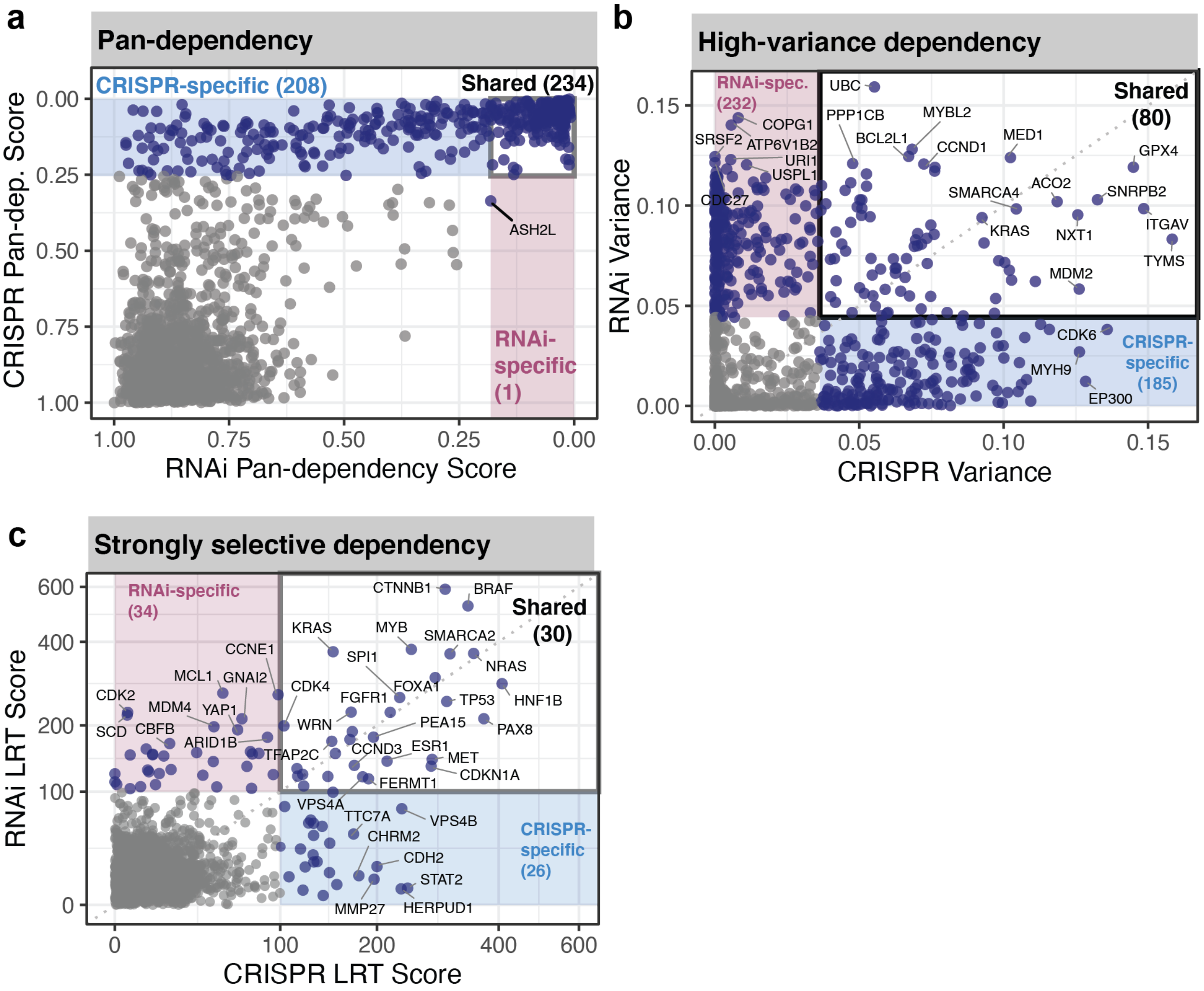


**Supplemental Fig. 12 | Shared and distinct pan-dependencies and selective dependencies**

**a**, Pan-dependency analysis (Methods) of high-confidence dependencies (N=1,703) for CRISPR (Avana) and RNAi (D2-Combined) datasets results in pan-dependency scores where scores closer to zero indicate that the 90th percentile of the gene effect distribution is closer to the strongest dependencies observed per cell line. Pan-dependencies distinct to either dataset are highlighted in colored boxes **b**, Variance in gene dependency (probability of dependency) for each high-confidence dependency (N=1,703) using matched CRISPR and RNAi datasets. Variance thresholds were calculated per dataset based on the variance of the 99th percentile of 643 non-expressed genes. **c**, Strongly selective dependencies (LRT > 100) are compared between CRISPR and RNAi datasets.

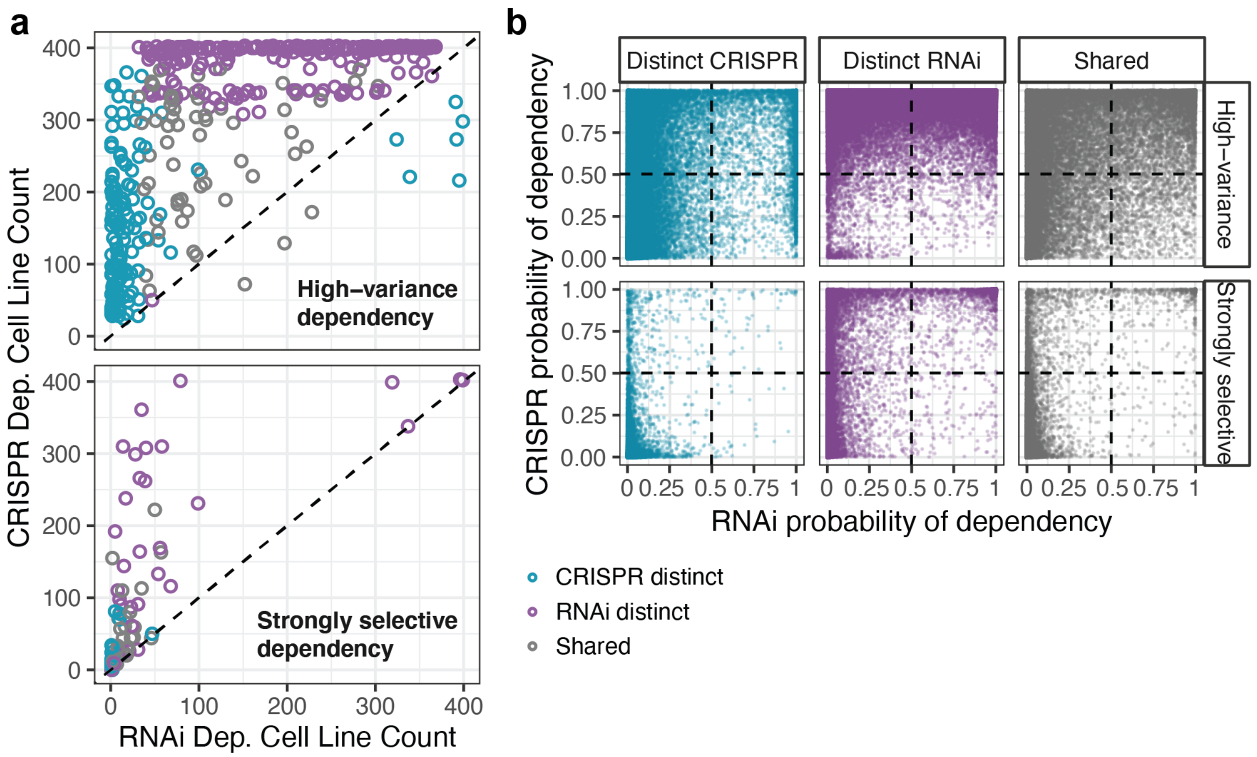


**Supplemental Fig 13 | Individual cell line dependencies detected using RNAi are supported using CRISPR, irrespective of whether selectivity patterns differ between perturbation types**

**a**, Number of dependent cell lines (probability of dependency > 0.5) per gene for the union of all high-confidence selective dependencies (high-variance, strongly selective) identified using CRISPR or RNAi datasets. Genes are colored according to whether the gene was classified as high-variance (top) or strongly selective (bottom) by only a single dataset (CRISPR distinct, RNAi distinct) or by both datasets (shared). **b**, Probability of dependency per cell line for all genes defined in part *a*. RNAi dependencies that are confirmed using CRISPR are in the upper right quadrant, whereas RNAi dependencies not confirmed using CRISPR are in the lower right quadrant.

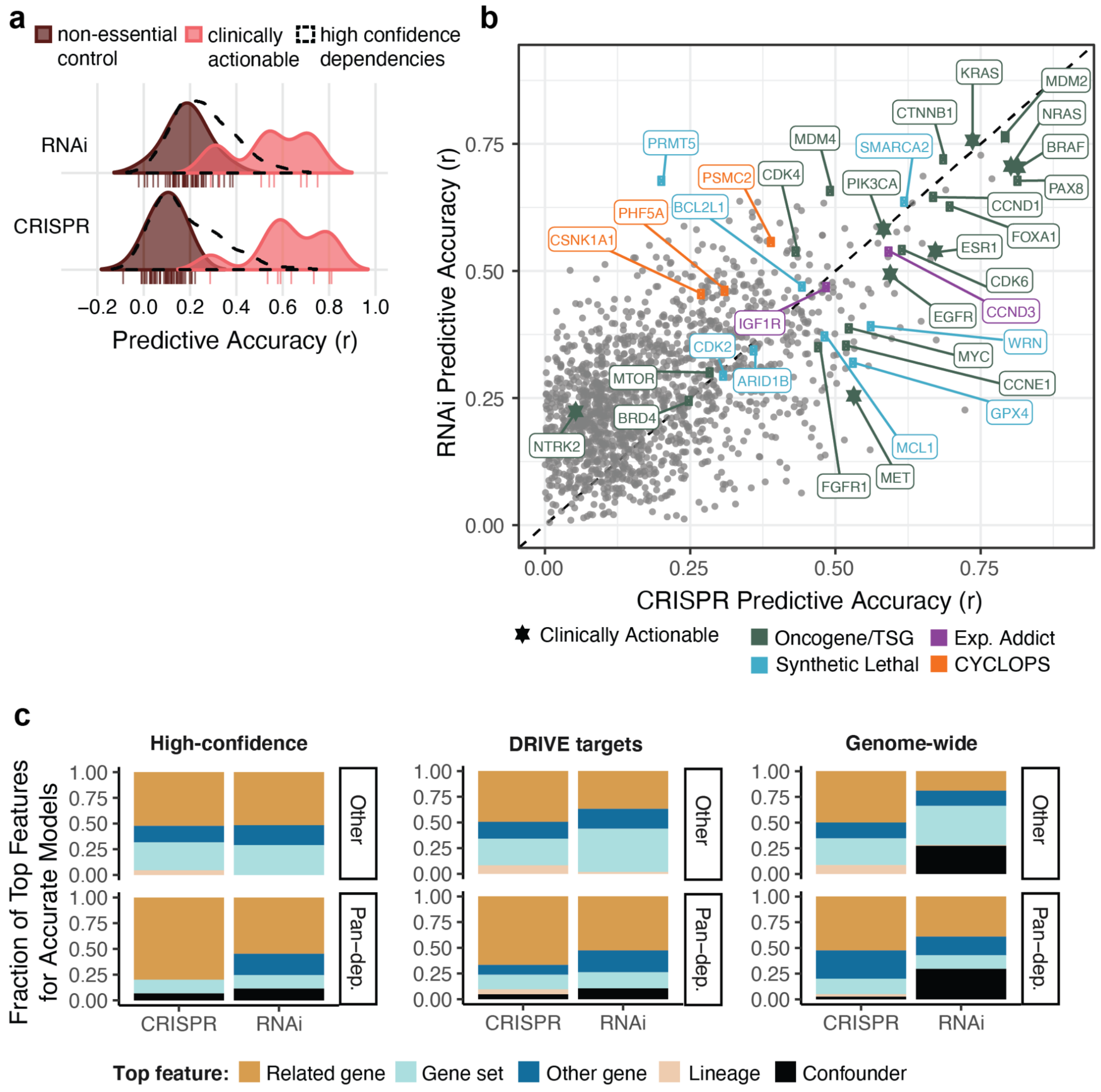


**Supplemental Fig. 14 | Benchmarking predictive accuracy and feature importance**

**a**, Multivariate regression of CRISPR or RNAi gene effect profiles using predictive features derived from multi-omics datasets and cell line annotations. Predictive accuracy is the correlation coefficient between measured and predicted gene effect values. Distributions represent non-essential (N=50), clinically actionable precision oncology targets with at least 5 dependent cell lines (N=7), and all high-confidence dependencies (N=1,701). **b**, Predictive accuracy annotated by clinically actionable precision oncology targets[6], oncogene and tumor suppressor genes (TSG)[6], and a curated list of established expression addictions, synthetic lethals, and CYCLOPS genes. **c**, The top predictive feature for each gene dependency profile, as determined by Gini feature importance, is annotated with its relationship to the gene target. Gene features are any CCLE multi-omics measurements of a single protein-coding gene. Related features are gene features that have prior information (CORUM, InWeb PPI, Ensembl

Paralogs) suggesting an association between the predictive and dependent genes. Confounding features represent screen quality or other technical aspects of the experiments. Gene set features are computed from RNAseq on a per cell line basis. Genes are faceted vertically by whether they are classified as CRISPR pan-dependencies and horizontally by whether they are included in the high-confidence dependencies, DRIVE targets, or genome-wide analysis.

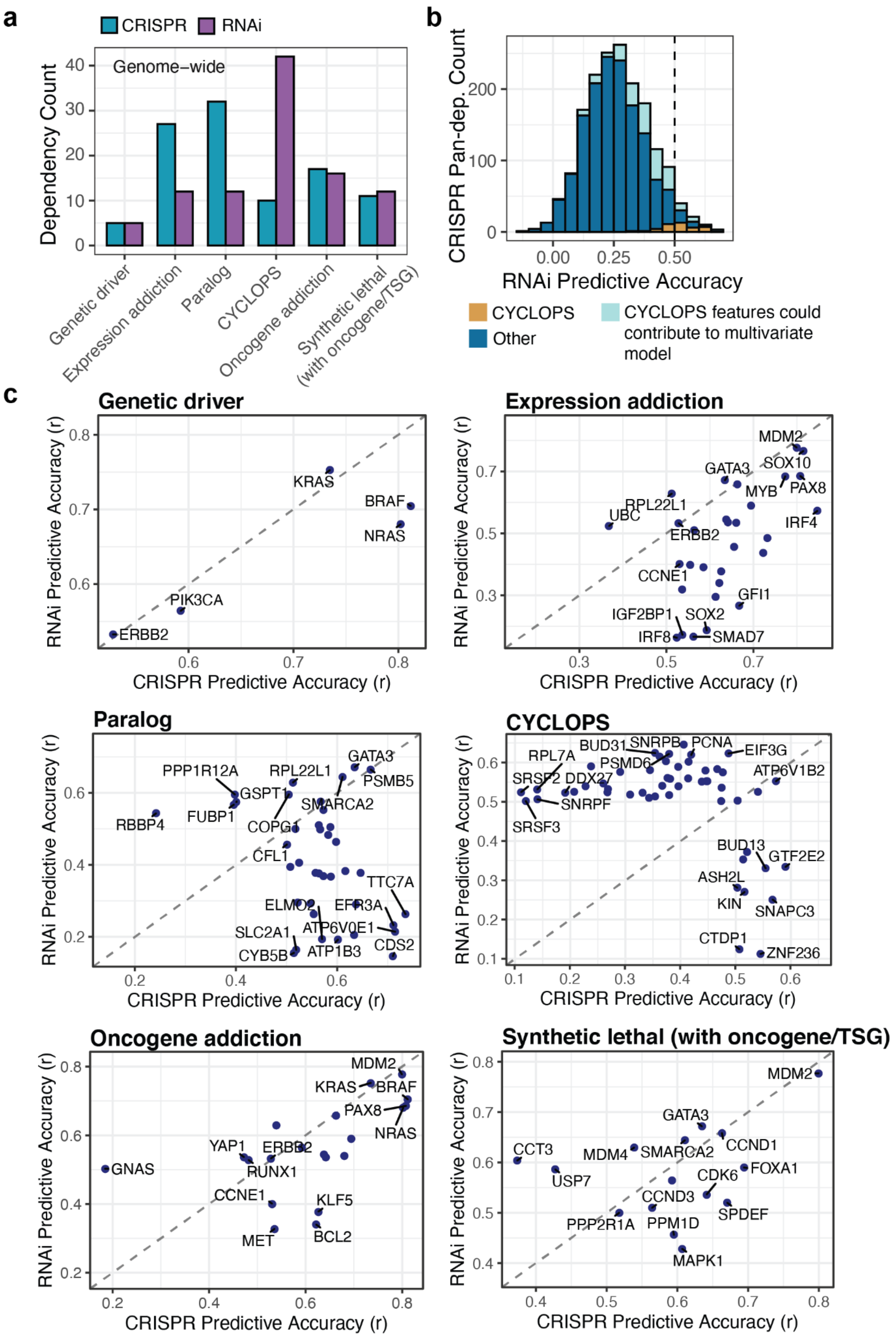


**Supplemental Fig. 15 | Genome-wide comparison of biomarker-dependency classes**

**a**, Number of predictive models (out of 15,221 genes shared between CRISPR and RNAi datasets) with predictive accuracy of at least 0.5, and predictive features which match the gene relationship, data type, and correlation direction defined by the biomarker classes. Details of biomarker class definitions can be found in Methods. **b**, Pan-dependencies detected using CRISPR are binned based on the predictive accuracy (x-axis) of the RNAi data for these genes. The colors represent the relationship between the RNAi gene effect profile and predictive features. Strong positive correlations (Pearson > 0.5) to between RNAi gene effect and copy number of the same gene are indicated as CYCLOPS. As an estimate of the number of genes that might have better predictive accuracy using RNAi due to some of the features in the multivariate model being CYCLOPS-related, we also annotated cases where the top multivariate predictor was copy number of a gene on the same chromosome arm (positive correlation) or expression of the perturbed gene target (positive correlation and left skewed gene effect profile). Genes without any clear relationship to copy number are labeled as ‘Other’. **c**, Genes included in each scatter plot are identified as the corresponding biomarker class using either perturbation type. The predictive accuracy shown is from the most accurate CRISPR or RNAi model for each gene dependency.

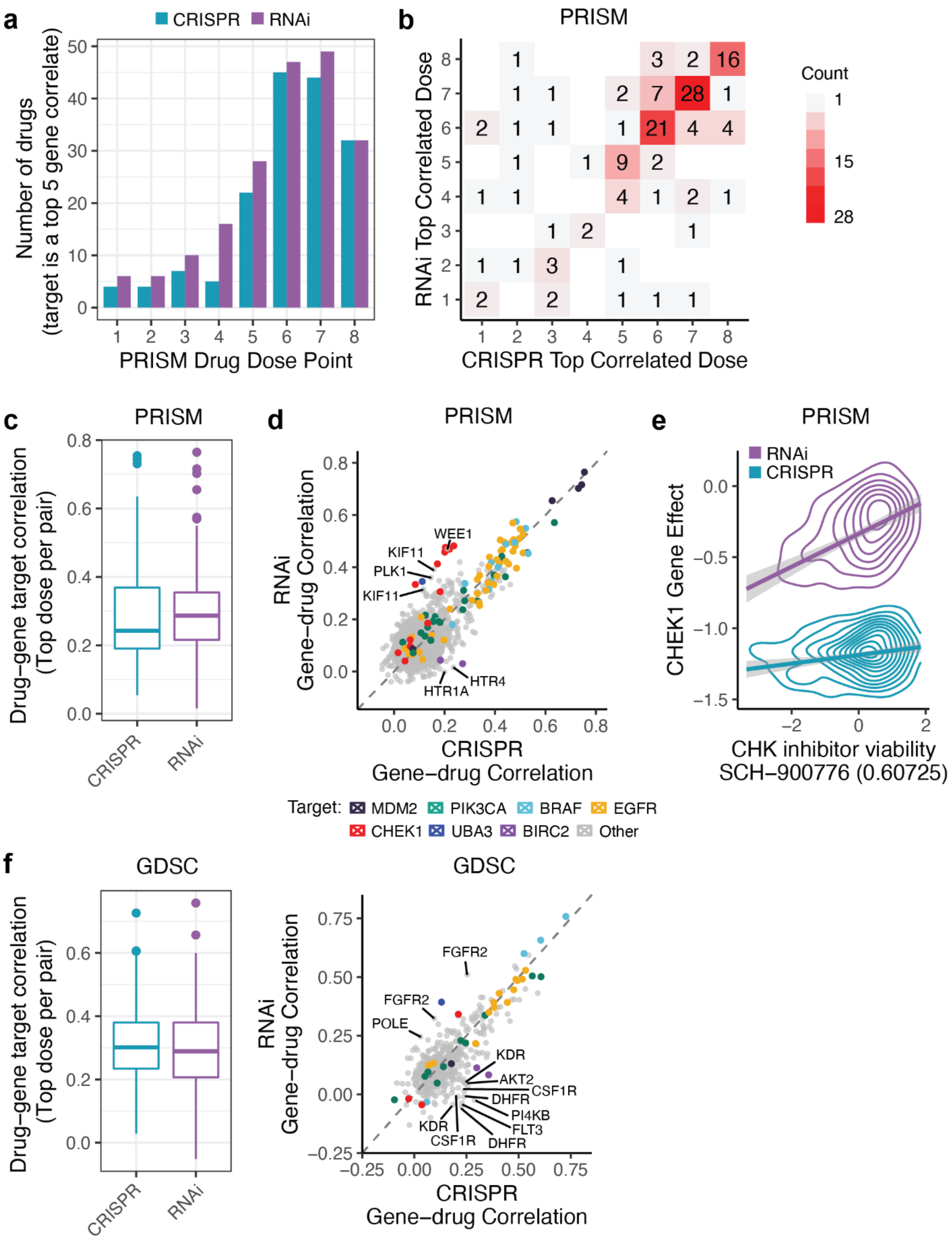


**Supplemental Fig. 16 | PRISM and GDSC drug-gene target associations**

**a,** Number of PRISM drug sensitivity profiles that have an annotated target within the top 5 correlated gene dependencies per drug dose. There are 558 drugs included that use a standard 8-point concentration range. **b,** Maximum correlated PRISM drug dose for each annotated drug-gene target pair (N=136) using CRISPR compared to RNAi. Drug-gene target pairs are included if the target is among the drug's top 5 gene correlates using either CRISPR or RNAi. Colors indicate the number of unique pairs in each bin. **c**, Pearson correlation of PRISM drugs and annotated targets (N=231) as in part *b* except without removing pairs for non-standard concentration ranges. **d**, Correlation of each PRISM drug with its annotated gene targets in the CRISPR and RNAi datasets (701 drugs, 304 gene targets). Multiple drugs targeting the same gene are colored according to the gene target. **e**, Density (2D) of PRISM CHK inhibitor (SCH-900776) viability at -0.60725 $\log_{10}$ concentration (M) and CRISPR or RNAi gene effects of its annotated target gene CHEK1. Data is smoothed using linear models with 95% confidence intervals. **f**, Pearson correlation of GDSC drugs and annotated targets (N=113) as was done with PRISM in part *c.* **g**, Correlation of each GDSC drug with its annotated gene targets in the CRISPR and RNAi datasets (214 drugs, 161 gene targets). Multiple drugs targeting the same gene are colored according to the gene target using the same color scheme as part *d*.

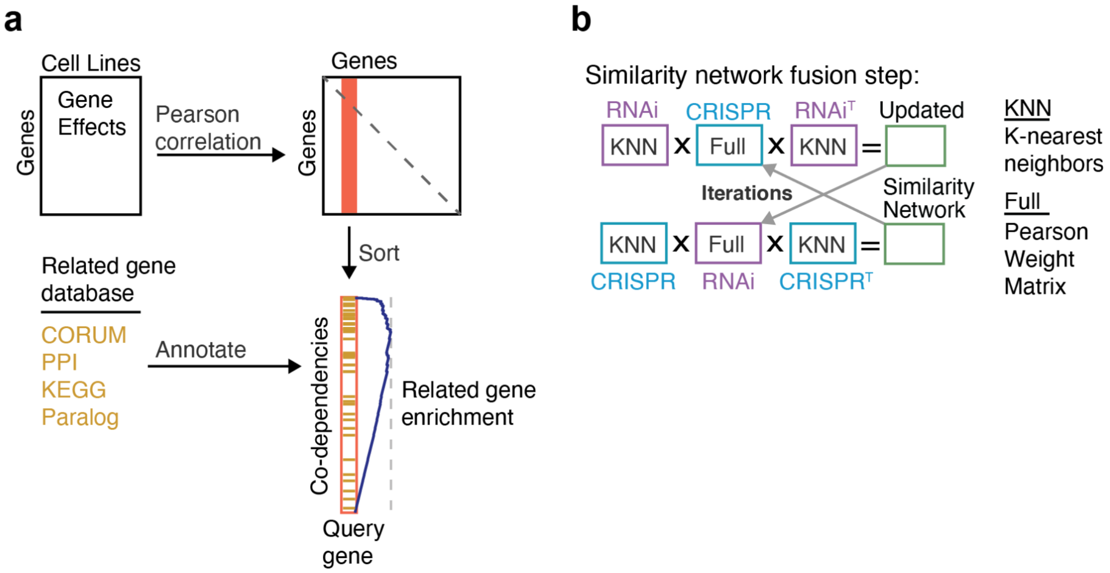


**Supplemental Fig. 17 | Co-dependency methods**

**a**, Approach to testing co-dependencies for recovery of prior information. **b,** Method of integrating CRISPR and RNAi into a single co-dependency network using Similarity Network Fusion (SNF).

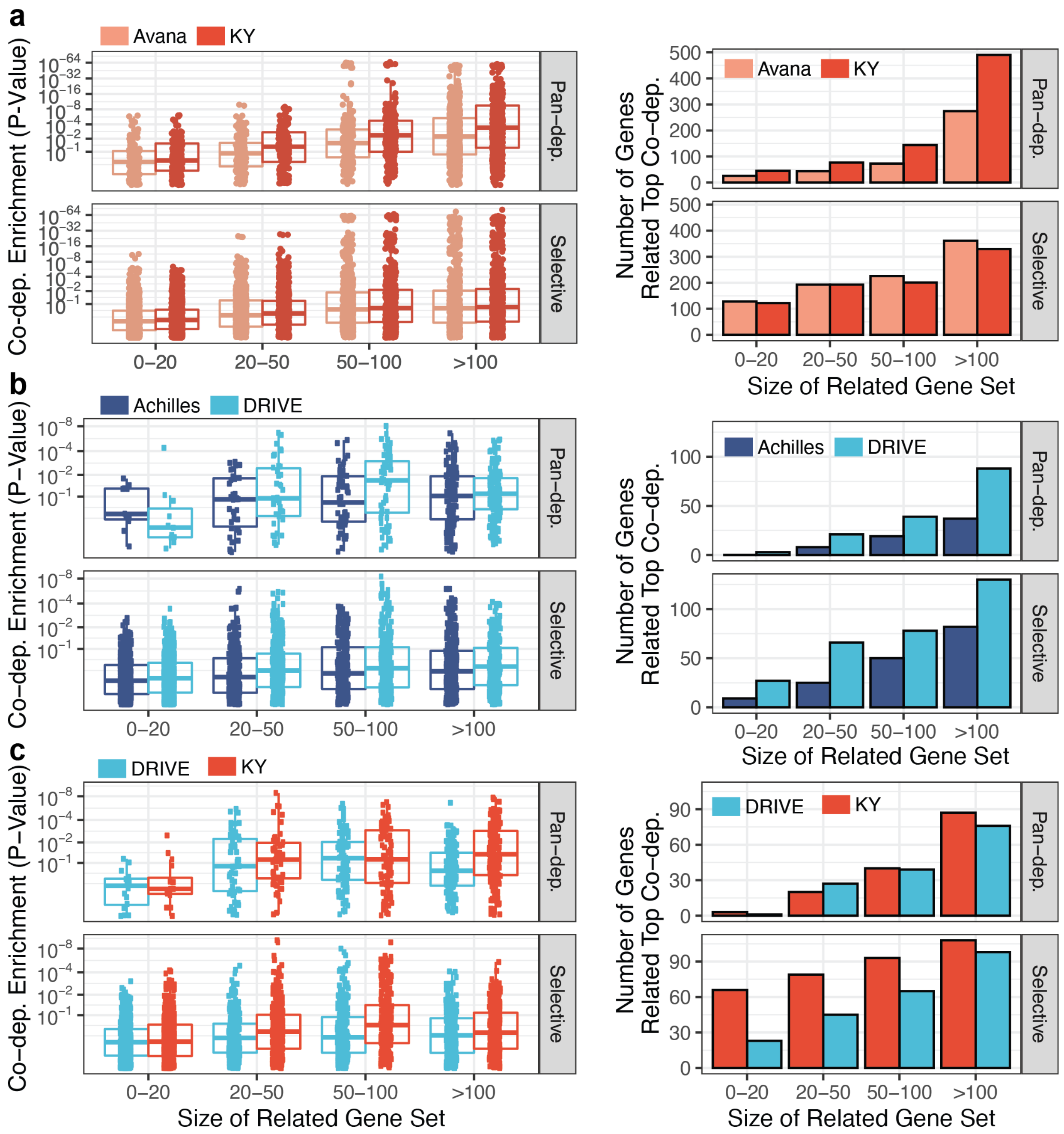


**Supplemental Fig. 18 | Recovering prior information from co-dependencies of each dataset**

**a**, A co-dependency network is constructed for each gene effect dataset (Avana, KY) by computing the Pearson correlation between all pairs of genes within the dataset (5,683 shared genes with at least 3 dependent cell lines in either dataset, 186 shared cell lines). The top correlates (co-dependencies) are queried for each gene and annotated by whether the co-dependent genes have been previously shown to be related (CORUM, PPI, KEGG, paralogs) to the query gene. The number of related gene priors can differ depending on the query gene (x-axis). Query genes that are classified as pan-dependencies in both datasets (N=1,314) are faceted separately (top). Enrichment of co-dependencies for related gene priors is tested for each query gene using a Kolmogorov-Smirnov test (left). The barplot on the right shows the number of query genes for which the top co-dependency is a related gene. **b**, Same as part (a) except networks are constructed using the 1,936 total genes, 226 cell lines, and 246 pan-dependencies shared between Achilles and DRIVE datasets. **c**, Same as part (a) except networks are constructed using the 1,880 total genes, 120 cell lines, and 337 pan-dependencies shared between DRIVE and KY datasets.

**a**

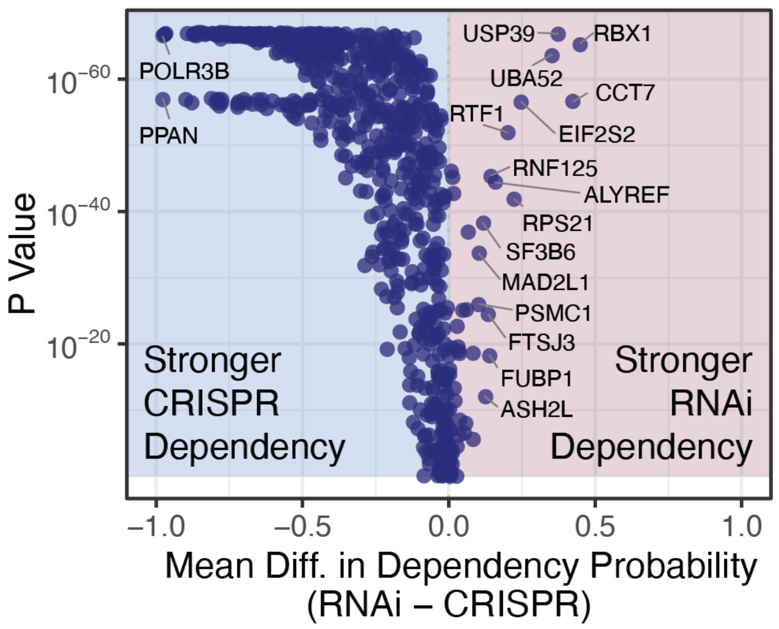


**Supplemental Fig. 19 | Comparison of mean gene effect using CRISPR and RNAi**

**a**, Wilcoxon test between RNAi and CRISPR dependency scores (probability of dependency) for each high confidence dependency gene that is a dependency in at least one cell line (N=800).
